# Protein catabolites as blood-based biomarkers of aging physiology: Findings from the Dog Aging Project

**DOI:** 10.1101/2024.10.17.618956

**Authors:** Benjamin R. Harrison, Maria Partida-Aguilar, Abbey Marye, Danijel Djukovic, Mandy Kauffman, Matthew D. Dunbar, Blaise L. Mariner, Brianah M. McCoy, Yadid M. Algavi, Efrat Muller, Shiri Baum, Tal Bamberger, Dan Raftery, Kate E. Creevy, Dog Aging Project Consortium, Anne Avery, Elhanan Borenstein, Noah Snyder-Mackler, Daniel E. Promislow

## Abstract

Our understanding of age-related physiology and metabolism has grown through the study of systems biology, including transcriptomics, single-cell analysis, proteomics and metabolomics. Studies in lab organisms in controlled environments, while powerful and complex, fall short of capturing the breadth of genetic and environmental variation in nature. Thus, there is now a major effort in geroscience to identify aging biomarkers and to develop aging interventions that might be applied across the diversity of humans and other free-living species. To meet this challenge, the Dog Aging Project (DAP) is designed to identify cross-sectional and longitudinal patterns of aging in complex systems, and how these are shaped by the diversity of genetic and environmental variation among companion dogs. Here we surveyed the plasma metabolome from the first year of sampling of the Precision Cohort of the DAP. By incorporating extensive metadata and whole genome sequencing information, we were able to overcome the limitations inherent in breed-based estimates of genetic and physiological effects, and to probe the physiological and dietary basis of the age-related metabolome. We identified a significant effect of age on approximately 40% of measured metabolites. Among other insights, we discovered a potentially novel biomarker of age in the post-translationally modified amino acids (ptmAAs). The ptmAAs, which can only be generated by protein hydrolysis, covaried both with age and with other biomarkers of amino acid metabolism, and in a way that was robust to diet. Clinical measures of kidney function mediated about half of the higher ptmAA levels in older dogs. This work identifies ptmAAs as robust indicators of age in dogs, and points to kidney function as a physiological mediator of age-associated variation in the plasma metabolome.

## Introduction

Lab-based studies on the biology of aging have led to major advances over the last several decades (Fontana et al., 2010; Kaeberlein et al., 2015; López-Otín et al., 2023). However, it is not clear how lab discoveries apply to aging in the real world, where variation in genotype, environment and their interaction, all present major challenges to translational geroscience (Partridge et al., 2018). More recently, modern molecular tools have made it possible to identify biomarkers for age, morbidity and mortality through the study of -omic domains, including the epigenome, transcriptome, metabolome, microbiome, and proteome (López-Otín et al., 2023). Among the -omic domains, here we focused on the metabolome, the collection of small molecules that make up the structural and functional building blocks of cells. Targeted metabolome profiles typically consist of measures of one to several hundred features. The metabolome integrates variation in vast numbers of environmental and genetic factors, whose effects converge onto a relatively small number of metabolomic endophenotypes (Panyard et al., 2022; Patti et al., 2012). The metabolome may thus reflect important axes of metabolic and physiological variation that underlie traits as complex as aging in nature.

One goal of gerontological research is to understand the causes and consequences of aging in humans. There are now many systems-level studies of age and aging in human populations (e.g. (Hannum et al., 2013; Horvath, 2013; Lehallier et al., 2019; Peters et al., 2015; Wilmanski et al., 2021)). There are a few limitations common to human studies. First, with few exceptions (e.g. (Kuo et al., 2022; van den Berg et al., 2023; Zhang et al., 2024)), studies of humans are carried out using cross-sectional designs, which introduce selection and survivor biases among other challenges (Nelson et al., 2020). The long average lifespan of humans introduces yet another challenge. In studies of middle-age or older humans, meaningful follow-up periods to assess either mortality risk, or longevity, of biomarkers exceed 10 years and may require 20+ years to achieve statistical power sufficient to identify the majority of biomarkers that portend future risk (Tessier et al., 2024). We need aging models in species with shorter lifespans and that parallel the complexity of the human environment and genetic variation. Into this space leaps the companion dog.

The companion dog has much to teach us about healthy aging and its associations with genetics and the environment (Creevy et al., 2016). Dogs vary tremendously, not only in size, shape and behavior, but also in their patterns of aging. Breed life expectancy can vary by a factor of more than two, from relatively short-lived giant breed dogs like Leonbergers and Mastiffs, to longer-lived small breeds, such as Pomeranians and Border Terriers (Yordy et al., 2020). Because dogs live with us, they experience the same local and regional environmental variation that we experience; they have a healthcare system as sophisticated as ours; and they have wide-ranging genetic variation. Most of these features are almost completely absent for laboratory models of aging. Moreover, the short lifespans of companion dogs relative to that of humans gives researchers a chance to see the impact their discoveries have on both dog and human health in their own lifetimes (Creevy et al., 2022).

In 2020, the Dog Aging Project (DAP) began enrolling tens of thousands of companion dogs in the United States to a long-term longitudinal study of normative aging (Creevy et al., 2022). The goal of DAP is to characterize the range of aging patterns in dogs, to discover the genetic and environmental factors that shape this variation, and to identify the mechanisms by which they do so. By combining information from owner-reported surveys, detailed demography, environmental data, veterinary electronic medical records (VEMR), ongoing studies of cognitive and behavioral traits, whole genome sequencing, clinical chemistry, and systems biology, and using epidemiological and public health research approaches, the DAP aims to identify how these factors influence dog heath and healthy aging (Creevy et al., 2022).

Though analysis of DAP data, we aim to shorten the timeframe for both biomarker discovery and analysis of normative aging, and to quicken the pace by which we test aging interventions in animals that mirror human genetic diversity and environmental complexity (Barnett et al., 2023; Creevy et al., 2022; Kaeberlein et al., 2016; Urfer et al., 2017). The few studies of age on the dog metabolome have revealed substantial differences between either the metabolome of cells cultured from young or old dogs (Brookes & Jimenez, 2021), or the blood plasma from dogs of a range of ages (Puurunen et al., 2022). Puurunen et al,. (2022) analyze plasma from over 2000 dogs and found age-associated variation in lipids, fatty acids and amino acids, amidst substantial variation due to diet, sex, and by breed.

Here we analyzed a panel of 137 aqueous metabolites measured in plasma collected from the Precision Cohort, which consists of a subset of DAP dogs recruited specifically for deep molecular profiling (Prescott et al., In review 2024). The data analyzed here were from 784 Precision Cohort dogs, representing a diversity of ages, during their first year of enrollment in the Cohort. We found that over a third of metabolites measured in dog plasma were associated with age, and we highlight three specific groups of age-associated metabolites, including acylcarnitines, indole derivatives, and post-translationally modified amino acids (ptmAA). These metabolites have been reported in studies of human age (Panyard et al., 2022), and here we investigated the physiological basis of age association with ptmAAs in particular. The only known source of free ptmAAs is the breakdown of protein, and we found additional evidence for protein catabolism within the metabolome. We found that clinical measures of kidney function at least partially mediate the age-associations of the ptmAAs. These results suggest that ptmAAs accumulate with age among dogs and may serve as a biomarker of aging physiology.

## Results

### Study Subject: The Dog Aging Project Precision Cohort

The Precision Cohort is a subcohort of the approximately 50,000 dogs that have so far been recruited into the Dog Aging Project. Beginning in January of 2021, 976 dogs were recruited into the Precision Cohort, representing a wide diversity of both dog age, genetics and geography. A summary of the cohort is shown in Figure 1. Here we analyzed metabolome data among 784 dogs from the first year of the Precision Cohort, which include 49% females and 51% males. Most dogs in the cohort were sterilized (87%). In designing the Precision Cohort within the overall DAP Pack, care was taken to recruit a cohort that reflects the full range of variation in American companion dogs in terms of geography, age, size, sex, sterilization status and purebred versus mixed breed status (Creevy et al., 2022). The Precision Cohort dogs with metabolome data at baseline resided in all but 1 of the 50 United States (**Figure 1A**), with an average of 15.7 dogs per state and 21%, 60% and 19% living in rural, suburban and urban environments, respectively.

**Figure 1.**
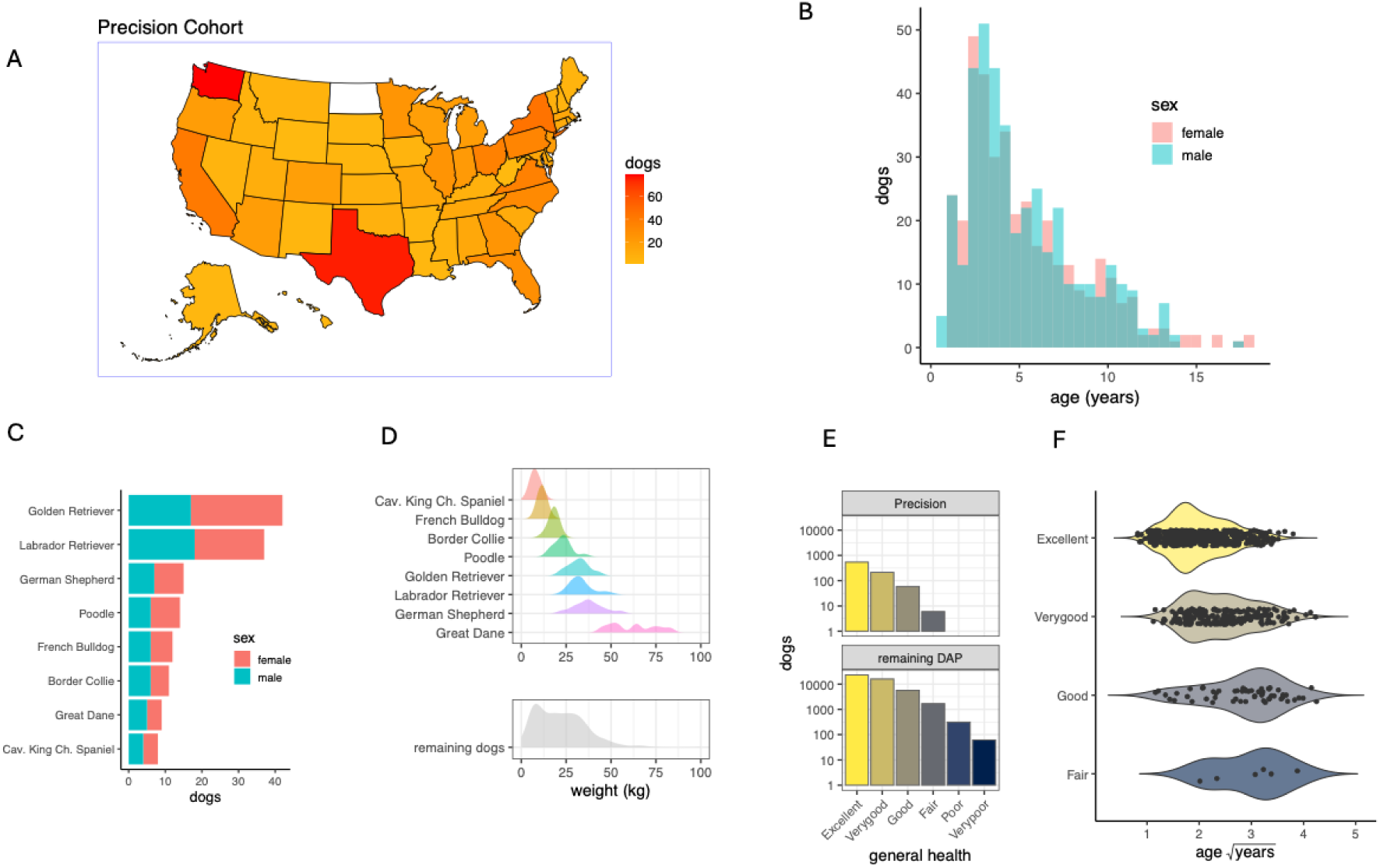
Demographic and health characteristics of the DAP Precision Cohort. (A)The geographic distribution of 784 dogs from the Precision Cohort. The number of dogs enrolled from each of the 50 United States is indicated by the color scale (range 1 to 79 dogs, white=0 dogs). (B) The age distribution by sex for the Precision Cohort (C) Based on the ancestry estimated from among 115,427 SNPs (Methods), 148 of the dogs consist of one of eight common breeds (those with at least eight dogs and at least 85% ancestry), and that also have representation from both sexes. The remaining 636 dogs were of either an under-represented breed, had ancestries from more than one breed, or did not include both sexes (Newfoundlands). (D) Dog weight at the time of blood collection for the most common breeds (upper panel), and for all other dogs (lower panel). (E) The number of dogs (note the log_10_ scale) at each general health category as given by owner reports. The upper panel summarizes the baseline Precision Cohort and the lower panel summarizes 47,444 dogs in the remainder of the DAP. Most Precision dogs (92%) were either in ‘Excellent’ or ‘Very good’ health, and no Precision dogs were categorized below ‘Fair’, whereas 0.8% of all DAP dogs were listed as being in ‘Poor’ or ‘Very Poor’ health. (F) Dogs reported to be in better general health were younger, on average, than dogs reported to be in worse health.

Among all dogs in the Precision Cohort, we identified evidence of ancestry from 110 different breeds (Sexton et al., *in prep* 2024, Methods). The maximum proportion of ancestry assigned to a single breed in a given dog ranged from 4.0% to 99.9%. To analyze the influence of breed on the metabolome, we chose dogs for which ancestry of the maximum breed exceeded 85% (Morrill et al., 2022), for breeds in which there were at least eight dogs in the cohort, and with representation from both sexes. This resulted in 148 dogs from eight breeds (**Figure 1C and 1D**). Each of the remaining 636 dogs were among a less common purebred group or had mixed ancestries contributed by a median of 5 breeds per dog (range of 2 to 38), and these 636 were treated as a single group (‘remaining dogs’) in breed-level analysis. Precision Cohort dogs were generally of good owner-reported health, with 92% (n=754) reported to have excellent or very good health status, and no dogs reported to have poor or very poor health (**Figure 1E and 1F**).

### Targeted plasma metabolomics

The metabolome data consisted of 137 aqueous metabolites on a targeted liquid chromatography-mass spectrometry (LC-MS) panel. The panel included amino acids and their derivatives, short-chain fatty acids and fatty esters, nucleotides, carbohydrates, organic phosphates and other metabolites.

### Multivariate metabolome analysis

We first investigated shared variation among metabolites in dog plasma using Principal Component Analysis (PCA). By comparing the distribution of data across each PC to that expected from random data, we found evidence that each of the first 23 PCs capture non-random variation (Tracy-Widom test, α<0.05); altogether, the first 23 PCs explain 63.3% of the variance in the metabolome (**Figure 2A**, Methods). Using Analysis of Covariance (ANCOVA), we estimated the variance of each PC that could be explained by the following variables: age, weight, sex, sterilization status, life stage (puppy, young, mature or senior), breed, the duration of fasting prior to blood collection, and 17 complete blood count (CBC) variables (Methods). These variables explained between 4.7% and 18.3% of the variance in each of the first 23 PCs (**Figure 2A**). Of all variables, age explained the most variance of any PC at 6.5% for PC4, and weight explained 5.6% of PC2 (**Figure 2A**). The CBC variables together explained up to 8.3% of the total variance among the first 23 PCs (**Figure S1B)**, while common breed, life stage, sex, sterilization status, and the duration of fasting each explained less than 5.0% of any of the PCs. While accounting for less than 5% of the variance, breed effects on the metabolome manifest across 5 of the first 23 PCs (P≤0.05, **Figure 2A**).

**Figure 2.**
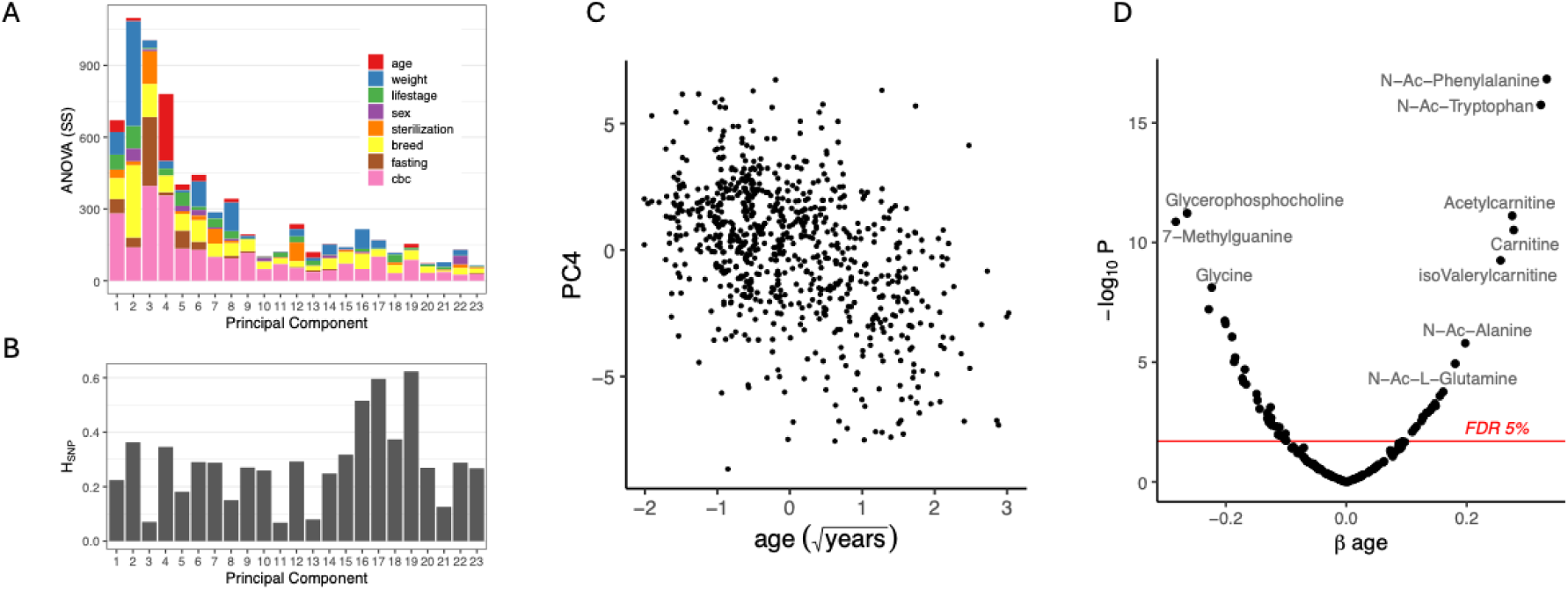
The age-associated dog plasma metabolome. (A) ANCOVA sum of squares (SS) among covariates (Methods) within the first 23 principal components (PC). The residual SS is not shown (see Supplementary Figure S1). (B) The proportion of variation in each PC that can be explained by the genetic relatedness among all dogs (H_SNP_, Methods). Note that the SS in (A) represents the total variance among the metabolome regardless of the PC, whereas in (B) H_SNP_ estimates the proportion of variance within each PC that was explained by relatedness. (C) PC4 associates strongly with age (square root-transformed years), Spearman’s *ρ*=-0.42, P<2.2×10^−16^. (D) The significance (−log_10_(P)) over the effect of age (β_age_) fit in a linear mixed model controlling for dog weight, sex, sterilization status, the duration of fasting prior to blood collection, CBC, and relatedness among the dogs (Methods). The FDR threshold of α = 0.05 is shown in red and representative metabolites are labeled.

**Figure S1.**
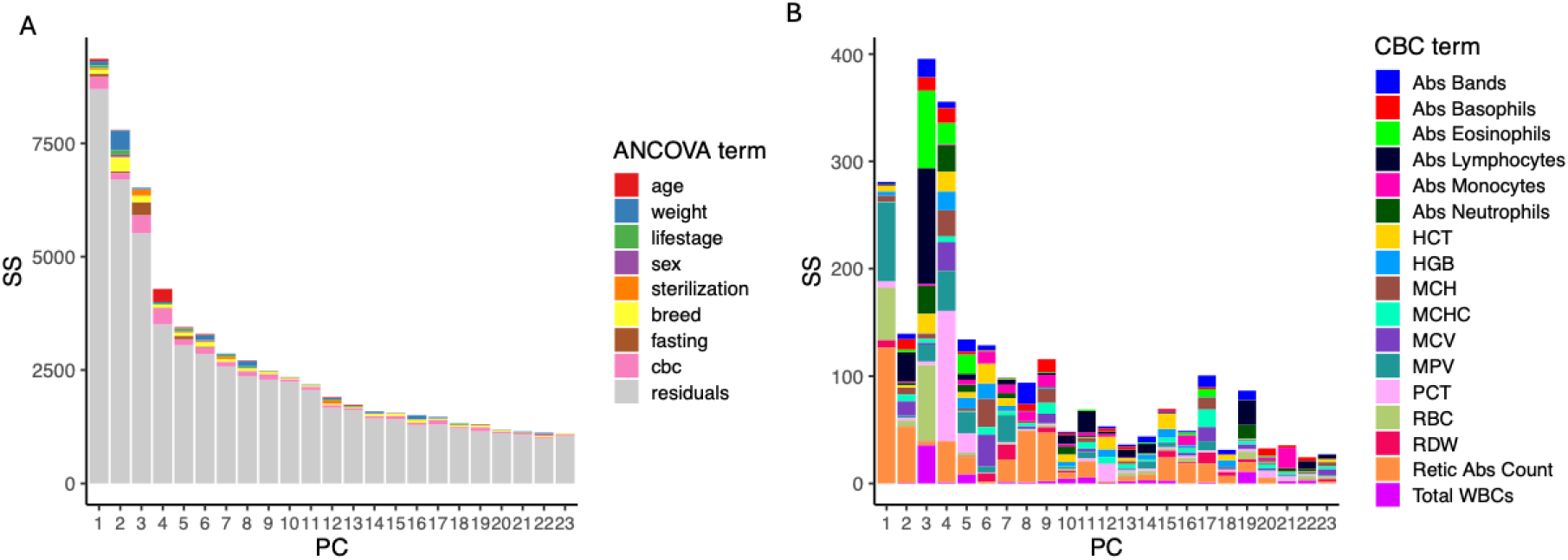
The multivariate dog plasma metabolome. (A) ANOVA sum of squares (SS) among the covariates (term) within each of the first 23 principal components (PC) of the plasma metabolome. The residual SS not accounted for by the terms is shown in gray. In (A) the SS from the 17 CBC traits are combined and indicated by the term ‘CBC’ (pink). (B) The SS for each of the 17 CBC traits (CBC term) across the first 23 PCs of the metabolome.

**Figure S2.**
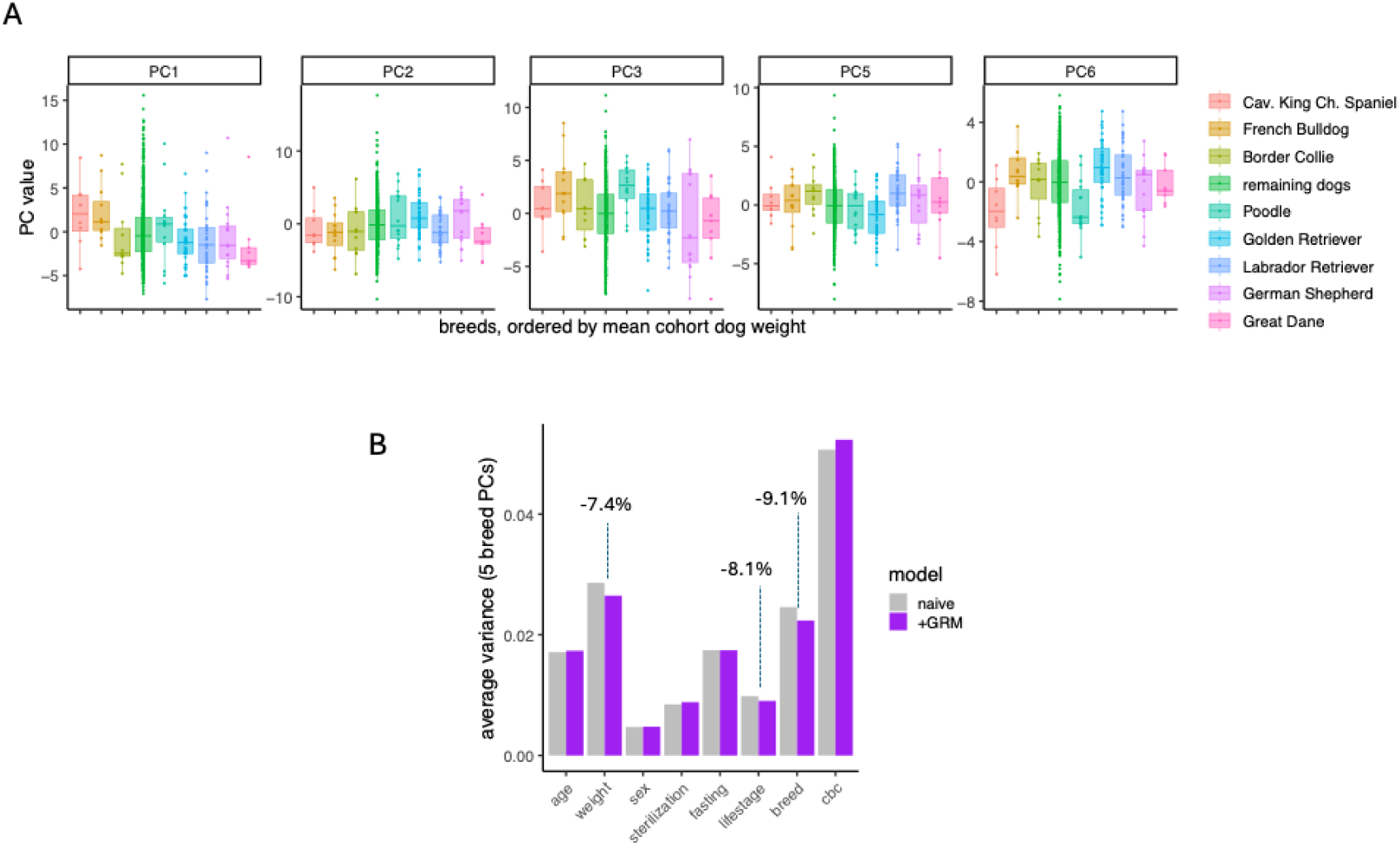
Breed has complex effects on the metabolome, partly accounted for by relatedness. (A) The 5 principal components (PCs) with effects of the 8 common breeds (ANCOVA P<0.05) plotted by breed, including all other remaining dogs. Within each plot, breeds are ordered by the mean weight of the dogs in each breed (n=8 to 44 cohort dogs per breed). (B) The average variance among the 5 PCs in (A) that was accounted for by the fixed effects indicated on the x-axis (BLUEs, Methods) in models that either include a random effect of relatedness by including the gene relatedness matrix (+GMR), or not (naïve, Methods). The percent reduction in average variance is indicated for the three most affected terms.

Given the difficulties associated with breed assignment in a genetically diverse cohort and the availability of low-pass sequencing data, we chose to dissect the multivariate metabolome variation attributed to the eight common breeds using the available genetic data. Using linear mixed effects models, we measured the degree to which the effect of breed on the metabolome could be explained by genome-wide relatedness. We fit each of the first 23 PCs to the fixed effects of the above-mentioned covariates along with the genetic relatedness matrix (GRM, Methods), which was treated as a random effect. Adding relatedness to the model reduced the effect of breed by 9.1% on average among the 5 PCs with breed effects (**Figure S2B**), indicating that only a modest proportion of variation due to breed can be explained by genome-wide relatedness.

In this diverse cohort, breed level effects were only evaluated for the 19% of dogs in the Precision Cohort that were one of the common breeds. The remaining dogs were either of mixed breed, or of breeds without sufficient representation to confidently attribute breed-level effects. Thus, to extend the analysis of genetic effects among all dogs, we asked to what extent the finer-scale relatedness could explain metabolome variation. For each PC, the proportion of variance explained by the GRM is referred to as its SNP-heritability (H_SNP_, (Yang et al., 2011)). Among the first 23 PCs, H_SNP_ averaged 28.3% with a range of 6.8% to 62.3% (**Figure 2B**). Therefore, in contrast to any genetic effects that may be represented by breed designation among the common breeds, the genomic data could hold more explanatory potential. In a separate study of the genetic associations within the plasma metabolome, we found that up to ∼60% of the variation in individual metabolites in the Precision Cohort can be explained by genetic relatedness (Sohrab et al., *in prep* 2024). In the following analysis, we controlled for the relatedness in order to ensure that genetic relatedness does not bias the results in our investigation of the effects of age on the metabolome. Age, which ranged from 0.7 to 18.0 years among the Precision Cohort, explained a significant portion of the variance (*P*<0.05) in 10 of the first 18 PCs, and was highly associated with PC4, where it accounted for 6.6% of the variance (**Figure 2C**, ANCOVA, *F_1,748_, P<*2×10^−16^).

### Effects of Age on the Dog Plasma Metabolome

Controlling for the covariates and relatedness, the effect of age (*β*_age_) was statistically significant for 54 of 137 metabolites at a false discovery rate (FDR) of α≤0.05 (**Figure 2D**). No metabolites showed a significant age x weight interaction effect (FDR>0.05), and so that term was removed from the model. Among age-associated metabolites, we observed two enriched groups of metabolites—the carnitines and the free forms of ptmAAs. Five of six carnitines on the targeted panel were age-associated, four were more abundant in older dogs and one, γ-butyrobetaine, lower in older dogs. Of the 21 metabolites that increase with age, nine were either a carnitine derivative or a ptmAA (**Figure 2D, Table S1**).

### Post-translationally modified amino acids as a biomarker of age

Older dogs have higher abundance of four ptmAAs, including N-terminally acetylated (N-Ac) N-Ac-alanine, N-Ac-phenylalanine, N-Ac-tryptophan, and N-Ac-glutamine (**Table S1)**. The three ptmAAs reduced in the plasma of older dogs compared to younger dogs were hydroxyproline, dimethylarginine, and N-Ac-aspartate (**Table S1**). The unmodified forms of each of the ptmAAs were also measured, and four of the unmodified amino acids were also age associated (**Table S1**). Age effects are prominent among the 12 measured ptmAAs (**Figure 3A**) We tested the hypothesis that the association between ptmAAs and age might be related to age-association of the corresponding unmodified version of each of the ptmAAs. The *β*_age_ of ptmAAs was not correlated with the *β*_age_ of the unmodified amino acids (linear regression *r^2^=*0.09, *P=*0.19). Similarly, of the 12 ptmAAs, the seven that were age associated did not correspond to which of the 12 corresponding unmodified amino acids were (Fisher’s exact test, odds ratio=0.24*, P*=0.5).

**Figure 3.**
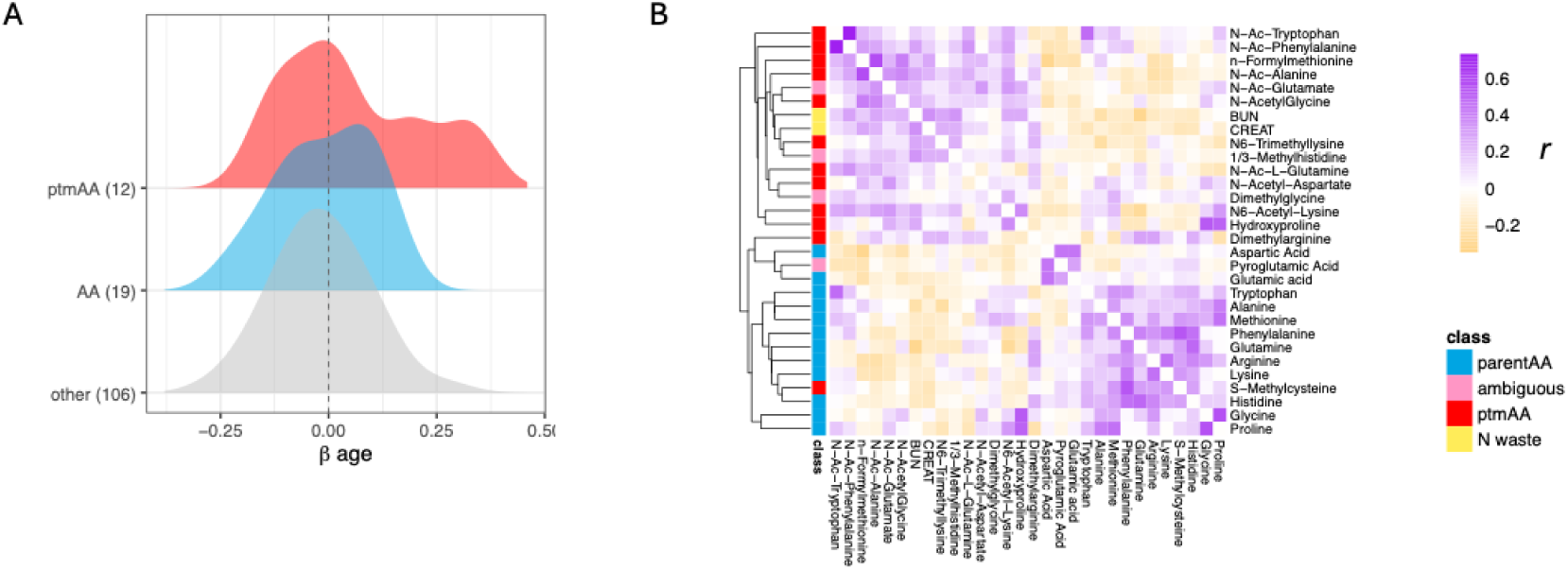
Post-translationally-modified amino acids associate with age and co-vary in plasma. (A) The distribution of the effect of age (β_age_) fit to each metabolite in a linear mixed model controlling for covariates (Methods). The distribution of β_age_ among the 12 quantified post-translationally-modified amino acids (ptmAA, red), the 19 quantified unmodified amino acids (AA, blue), and the 106 remaining metabolites of other classes (other, grey). (B) A heatmap of clustered pairwise correlation among the ptmAA, with the addition of four modified amino acids that are not necessarily port-translationally modified and may form either from protein hydrolysis, or from de novo synthesis (ambiguous), the unmodified amino acids (parent AA), and two nitrogenous waste products (N waste): creatinine (CREAT) and blood urea nitrogen (BUN). Values in the map are Pearson’s r from correlations calculated among the residuals of the full mixed model, and so are adjusted for age, relatedness and the other covariates. The diagonal was made white for clarity. The rows are annotated by metabolite class and a dendrogram of UPGMA clustering (Methods) is shown at left.

There are two alternative explanations for the difference in abundance of ptmAAs in young and old dogs. First, there might be broad changes with age in the influx or removal of ptmAAs from the blood, from a common source, such as might be caused by differences in protein catabolism, or the removal of its byproducts. Alternatively, there might be a variety of processes that could generate or remove individual ptmAAs in plasma, such as the turnover of particular endogenous proteins, the transport of specific ptmAAs from the gut to the bloodstream, or the degradation or excretion of select ptmAAs. While we do not rule out the latter possibility, we found empirical support for the former hypothesis. If protein catabolism broadly differs among young and old dogs, then, *a priori*, this should give rise to ptmAAs in rough stoichiometry to their abundance among digested proteins. Therefore, we would anticipate that, broadly, the abundance of ptmAAs would positively co-vary in plasma. Alternatively, if ptmAAs were acquired, synthesized or removed independently, without a common origin, their abundances would not be expected to correlate. We examined the correlation among metabolites after removing the effects of age and other covariates (Methods). By clustering the Pearson’s correlation among both the ptmAAs and their unmodified forms, we found that each metabolite covaried positively with between 2 and 16 other metabolites (FDR<0.05), and that the ptmAAs and unmodified AAs cluster separately (**Figure 3B**). The covariation among ptmAAs supports the hypothesis that there is a general age-related shift towards protein catabolism and/or diminished removal of its byproducts in dogs.

The primary physiological sources of amino acids from protein catabolism are from proteolysis of dietary protein by microbiota in the intestine, digestion of protein in muscle and other tissue, and proteolysis in the liver (Denton & Elvehjem, 1954; Levitt & Levitt, 2018). Amino acids originating either from intestinal hydrolysis of protein, or from tissue protein digestion, circulate to the liver via the portal vein where, if they are metabolized, generate nitrogenous waste in the forms of creatinine and urea. Creatinine was measured on the targeted panel and the blood samples also had standard clinical blood chemistry measured, which includes blood urea nitrogen (BUN) and serum creatinine. As expected by the protein catabolism hypothesis, the level of serum creatinine and of BUN correlate positively with the abundance of 11 and nine of the 13 ptmAAs, respectively (mean *r* = 0.125 and 0.165, P<0.05), and not with any of the unmodified AAs (mean *r*=−0.101 and −0.109, P>0.05 **Figure 3B**).

### Testing putative sources of post-translationally modified amino acids

The age-association of ptmAAs could be caused by age-related changes in metabolism, including the catabolism of either dietary protein, or endogenous sources of protein like the digestion of tissue or cells. Alternatively, age may be accompanied by changes to the rates of clearance of ptmAAs from the blood. We evaluated variation in diet as a potential driver of ptmAA in plasma. Drawing on survey responses from 761 dog owners, we found these dogs vary substantially in their primary diet category from the most popular, dry kibble, to raw, canned and freeze-dried foods, including representation from both commercial and home-prepared diets (**Figure S3A**). We tested each diet type for associations with plasma metabolites using dry kibble, the primary diet of 86% of P1 dogs, as the reference diet (Methods). There were up to 30 metabolites associated with at least one diet type, with the effects of home-prepared (raw or cooked), and raw commercial diets being similar (**Figure S3B**). Of the age-associated ptmAAs, only N-Ac-aspartate was affected by diet, being lower in abundance in plasma of dogs primarily eating commercial raw diets (refrigerated or frozen raw) in comparison to dry kibble (β=-0.49, FDR=0.023, **Figure S3C**). Additionally, the ptmAA S-methylcysteine, which was not age-associated, was higher in dogs eating commercial raw and home cooked diets than those eating kibble (**Figure S3C**).

**Figure S3.**
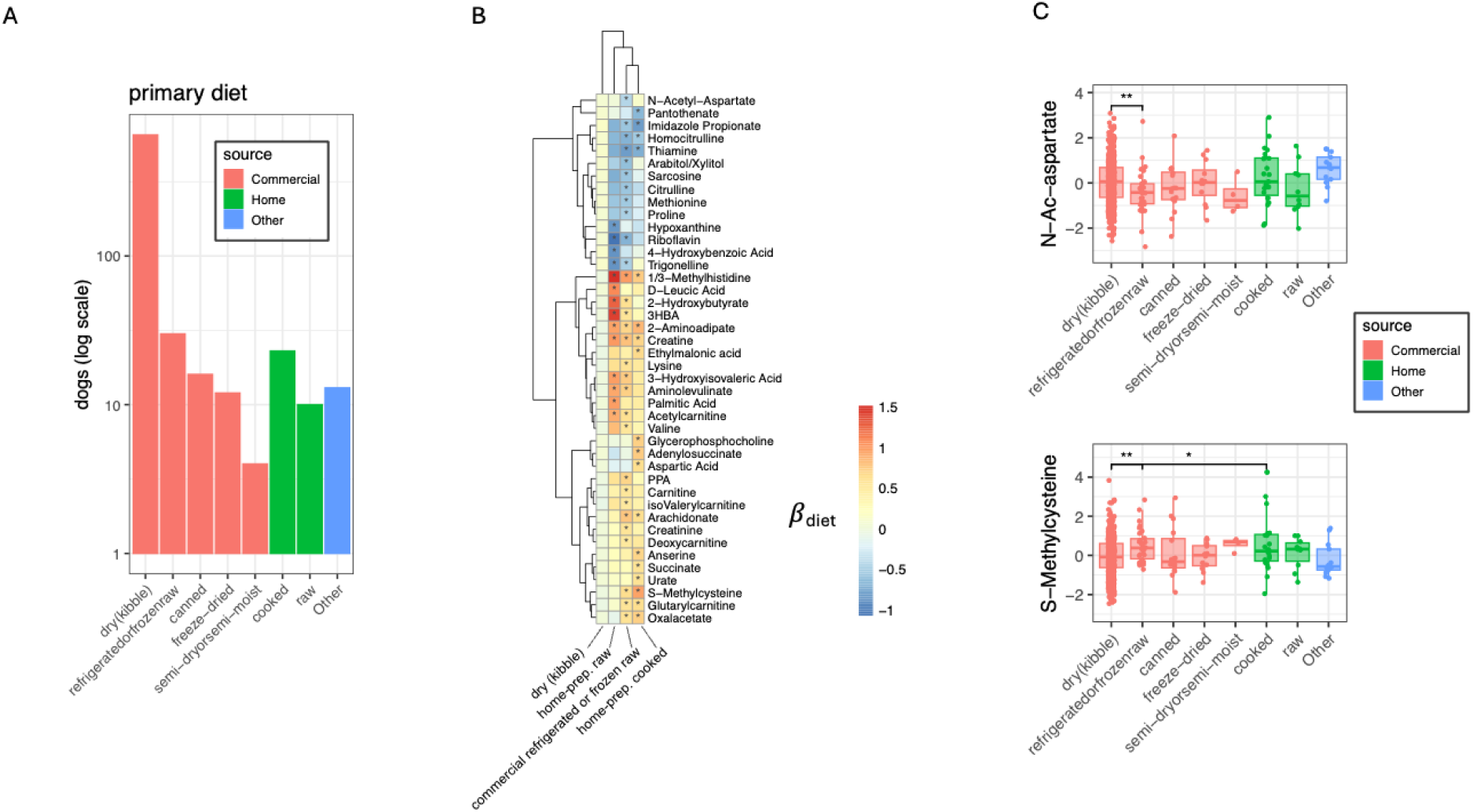
Primary diet composition does not explain variation in post-translationally modified amino acids in plasma. (A) The distribution of the primary diet type among owner survey responses of 761 dogs, with counts plotted on a log scale. Diet components were divided into those commercially sourced (Commercial), home prepared (Home), or of some other type (Other). Non-responses were omitted, and so Other reflects diet components that do not fit one of the seven categories or where diet was not consistent. (B) A heatmap showing the effect of diet type on each metabolite (*β*_diet_) in comparison to dry kibble in a mixed model to control for age, weight, and other covariates (Methods). Alongside the metabolite values among the reference diet, only metabolites and diets that had at least one effect are shown (*β*_diet_≠0, FDR < 0.05, asterisks). (C) N-Ac-aspartate and S-methylcysteine were the only post-translationally modified amino acids to associate with any primary diet component (FDR<5%, asterisks).

As an additional means to identify effects of age while controlling for the effect of diet, we tested for effects of age only among the 653 dogs that eat kibble as their primary dietary source. Even with reduced statistical power due to smaller sample size, all four of the ptmAAs that were positively associated with age among all dogs, and two of the three negatively associated metabolites, hydroxyproline and dimethylarginine, remained age-associated (FDR<0.05). N-Ac-aspartate, which was negatively associated with age, was no longer age-associated among the dogs primarily eating kibble (FDR=0.14). While diet may influence the ptmAAs that were lower in older dogs compared to younger dogs, we found no evidence that variation in primary food type among the dogs explains the increased abundance of ptmAAs in the plasma of older dogs.

The kidneys play a key role in removing metabolomic byproducts to the urine, including the waste products of protein and amino acid catabolism. To test for effects of kidney function on plasma ptmAA, we evaluated several potential biomarkers. Serum creatinine is a common clinical metric used to estimate glomerular filtration rate (GFR, (Dahlem et al., 2017; Hokamp & Nabity, 2016; Yamaguchi et al., 2021)). Given that serum creatinine co-varies with many of the ptmAAs in plasma (**Figure 3B**), we tested the potential for creatinine to act as a mediator of the elevated ptmAAs in the plasma of older dogs (Methods). By comparing the effect of age on each metabolite, with and without the addition of creatinine to a full mixed model, we found evidence that the age association of six of the seven age-associated ptmAAs were substantially mediated by serum creatinine (**Figure 4A**, FDR<5%, Methods). Therefore, for the ptmAAs other than hydroxyproline, if we account for serum creatinine, which was inversely proportional to the GFR, we can account for between 45 and 68% of the effect of age.

**Figure 4.**
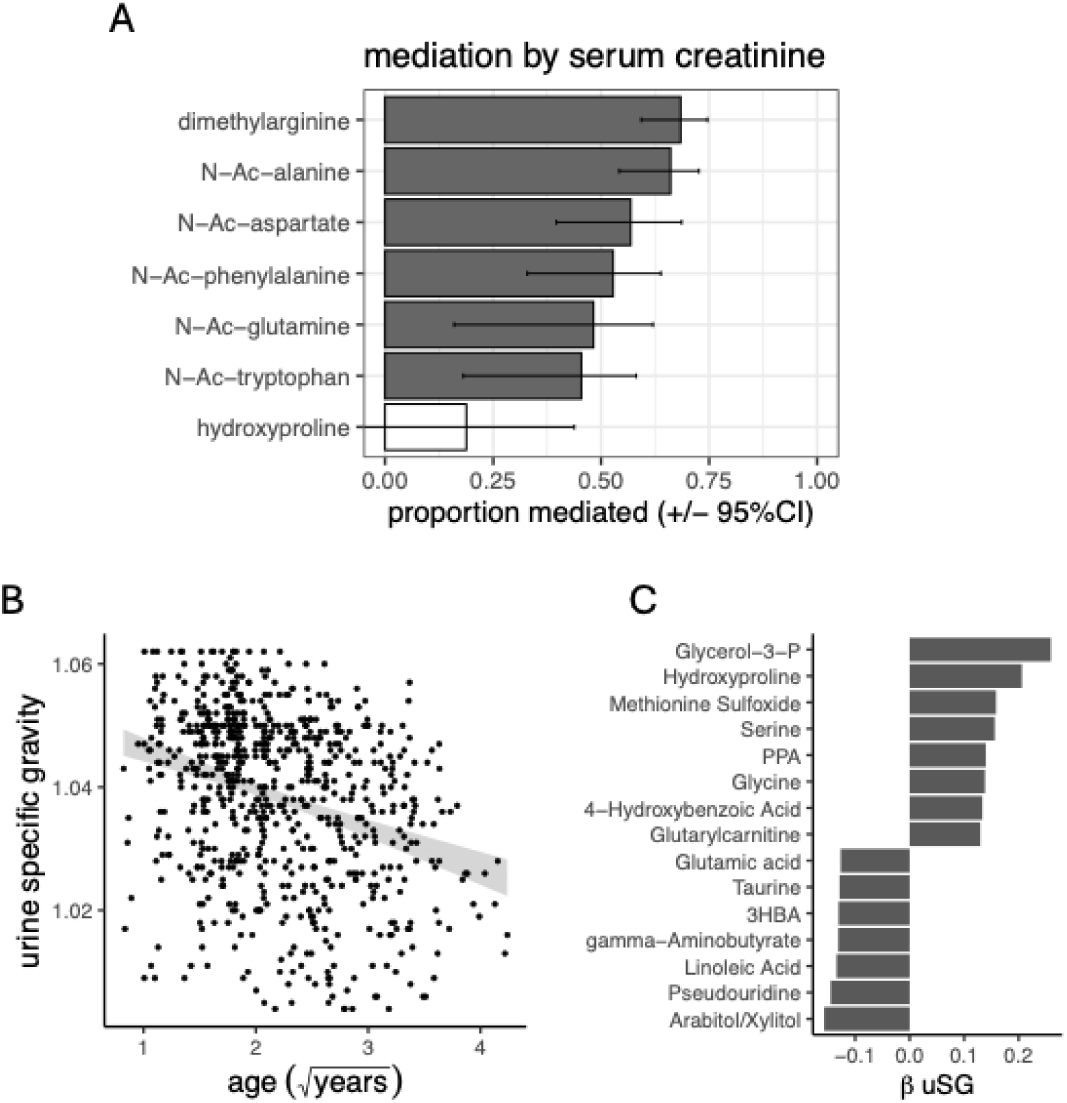
Indicators of kidney function partially explain the age-association of ptmAAs. (A) Of the 7 ptmAAs that associate with age, the proportion of the age effect on ptmAAs in a full mixed model that remains (proportion mediated) after the addition of serum creatinine to the model (error bars are 95% confidence intervals, Methods). Filled bars correspond to metabolites mediated by creatinine (ACME, FDR<5%). (B) Urine specific gravity (uSG) from clinical urinalysis associated with age among 741 dogs in a mixed model controlling for covariates (FDR = 1×10^−10^, Methods), gray shaded region indicates 95% confidence interval for ordinary least squares regression. (C) The effects of uSG (β uSG) on each of 15 metabolites while controlling for covariates (FDR<5%, Methods).

We sought alternative indicators of kidney function from clinical urinalysis of 741 of the Precision dogs. Among 10 urinalysis measures, both bilirubin and urine specific gravity (uSG) declined with age (FDR<5%, **Figure 4B**). We failed to find metabolites associated with bilirubin. However, there were 15 metabolites, including hydroxyproline, associated with uSG (**Figure 4C**, FDR<5%). No other ptmAA associated with uSG. Urine specific gravity was not associated with serum creatinine (linear mixed model, β_creatinine_, P=0.059), and therefore uSG could indicate kidney function that was independent of serum creatinine. While uSG was associated with hydroxyproline when controlling for all other covariates, uSG was not a mediator of age on hydroxyproline (FDR>23%).

## Discussion

We have surveyed the age-related plasma metabolome among the diverse Precision Cohort of the Dog Aging Project. Amidst the genetic and environmental complexity of the companion dog, we found metabolites associated with CBC/Chem variables, dog weight, sterilization status and the duration of fasting, as well as significant variation by breed, only some of which could be explained by finer-scale genetic relatedness. In controlling for these covariates we sought plasma biomarkers of age with higher translational potential. In doing so, we found substantial differences in the plasma metabolome of dogs by age, including almost 40% of metabolites. The age-associated metabolites were similar to those in humans, and include byproducts of protein catabolism. We then query the available data for the Precision Cohort and identify diet and biomarkers of kidney function as potential mediators of parts of the age-associated metabolome.

While most studies of dogs aimed at identifying biomarkers of age or health focus on dogs of a particular breed (Morelli et al., 2022; Qu et al., 2022), or on dogs living in the limited environment of a dog colony (Christie et al., 2009), the Precision Cohort of the DAP is similar to the large-scale cohort of Puurunen et al (2022), with the shared aim for a comprehensive representation of a wide range of dog genetic and environmental variation. Similar to Puurunen *et al*. (2022), we saw indications of genetic influences throughout the metabolome. Here we made use of genome-wide allelic variation to show that a substantial portion of the metabolome was explained by fine-scale relatedness among dogs, which points toward genetic influences on the metabolome that transcend breed level variation. For sake of comparison to the large cohort analysis of Puurunen *et al*. (2022), however, we first considered a breed-level analysis. The effects of breed that we report are less extensive than those in Puurunen *et al*. (2022). We evaluate breed level effects with caution, as others have demonstrated that traits perceived as breed-specific are often better explained by finer-scale genetic analysis (Morrill et al., 2022). We then used linear regression to decompose the metabolome variation associated with breed and found that only a modest portion of breed-level effects could be explained by genetic relatedness. Given the limitations of breed-level analysis in a diverse cohort, we focused our analysis of age effects in a more general model that simply controls for relatedness and other covariates.

Of all biological covariates considered here, age had the most substantial influence on any single principal component of the metabolome (**Figure 2**). However, we note several likely sources for bias in this study cohort, all of which lead us to reason that the baseline characterization of age-associations among the Precision Cohort may reflect healthy aging, rather than indicators of unhealthy aging (Nelson et al., 2020). A baseline cohort of any species, particularly one that recruits both younger and older participants, is inherently subject to survivor bias, where subjects in the study, particularly at later ages, can only represent the subset of individuals who have survived to that age (Anderson et al., 2011). Furthermore, an owner’s decision to enroll a dog can be influenced by their perception of whether their dog is a good fit for the study, creating a self-selection bias. For instance, owners may avoid enrolling dogs they perceive as “too young”, “too old”, or “too sick.” This mirrors the volunteer bias in human studies, where individuals with certain health status or demographics are often underrepresented (Fry et al., 2017a; Galea & Tracy, 2007). Last, bias can arise when participation is skewed toward those with more resources. Owners with more resources—time and money—to participate in a longitudinal study are more likely to enroll their dogs, leading to a sample that may not accurately reflect socioeconomic status in the broader dog-owning population (Fry et al., 2017b). While acknowledging these caveats, we discuss our results as indications of age-related variation in the plasma metabolome, and what it might indicate about the physiology of dogs as they age.

We used a mixed model framework to estimate age-association in the univariate plasma metabolome while simultaneously correcting for the covariates mentioned above. In doing so, we avoided the confounding influence of common variation among dogs with the aim of identifying age-associations that are more likely to translate to dogs generally. We found a significant effect of age in 39% of plasma metabolites (**Figure 2. Table S1**). We focus our discussion on three groups of metabolites—carnitines, indole-derived metabolites, and the ptmAAs and other products of protein catabolism—for their similarity to age-associated metabolites in humans, and with regard to the ptmAAs, because they provide novel clues to the physiological differences between young and old dogs. Each of these groups includes metabolites that are associated with age in human plasma, and in the following sections, we summarize these associations and their similarities to that in humans and discuss potential physiological processes that may lead to this variation.

### Age-associated plasma metabolites in dogs, and parallels in humans

**Acylcarnitines**: Among the most consistently age-associated plasma metabolites in humans and mice are those involved in fatty acid metabolism, including the fatty acids themselves, as well as carnitine and acylcarnitines (Darst et al., 2019; Houtkooper et al., 2011; Jarrell et al., 2020; Johnson et al., 2019; Lassen et al., 2023; Robinson et al., 2020; Sol et al., 2023; Tessier et al., 2024). In this study, we limited our analysis to the aqueous metabolome, which lacks hydrophilic lipids. However, of the six carnitines measured, four associated strongly with age, with all but one, γ-butyrobetaine, positively associated with age. Acylcarnitines and carnitine are required to shuttle fatty acids into mitochondria for β-oxidation, and several authors speculate that the rise in acetylcarnitine with age in humans could be due to reduced function of mitochondria (Jarrell et al., 2020; Lassen et al., 2023), or of the kidneys (Yamaguchi et al., 2021). γ-butyrobetaine, the only carnitine negatively associated with age, is interconverted with carnitine, which associated positively with age. The opposite relationship of carnitine and γ-butyrobetaine therefore could be due to the age-related change in balance between carnitine and γ-butyrobetaine. Overall, the increased abundance of acylcarnitines in plasma among older dogs was consistent with patterns seen in humans, although the cause is unclear.

**Tryptophan/Indol metabolites**: In contrast to many metabolites whose sources are ambiguous, indole-conjugated metabolites found in plasma are generated exclusively by the conversion of tryptophan by gut microbes (Sinha et al., 2024; Wikoff et al., 2009). Of the three indol-3 derivatives measured here, indole-3-propionate was lower in the plasma of older dogs, and indole-3-lactate was higher. Metabolites related to tryptophan and its indol derivatives are also associated with age in human plasma (Johnson et al., 2019; Lassen et al., 2023; Sol et al., 2023). Associations between the abundance of microbial-derived metabolites and dog age strongly suggests that gut microbial metabolism influences the age-related plasma metabolome.

The fecal microbiome of older dogs in the Precision Cohort is both less diverse and is more unique between older dogs than in younger dogs (Bamberger et al., *in prep* 2024). Both of these age-related trends are also observed in humans (Wilmanski et al., 2021). Furthermore, reduced diversity of the human microbiome associates with plasma metabolites, and with longevity (Wilmanski et al., 2019). That indol-derived metabolites were age-associated in dog plasma suggests a role for the microbiome in dog aging and its influence on the plasma metabolome. Future longitudinal multi-omic analysis could test for effects of tryptophan derivatives and the role of the microbiome in dog aging.

**Byproducts of Protein Catabolism:** ptmAAs are only known to form on polypeptides or, in the case of N-formyl-methionine, on methionine-charged tRNA prior to protein synthesis (Ree et al., 2018). Therefore, the only known source of these metabolites, in the free forms measured by LC-MS, is protein catabolism. We identified four additional age-associated modified amino acids—pyroglutamate, 1/3-methylhistidine, dimethylglycine, and N-Ac-glutamate. However, each of these can be generated either post-translationally or by modification of free amino acids, and so their source is ambiguous. Studies of plasma metabolites often include measures of the amino acids, and in both dogs and humans, amino acid concentrations are regularly found associated with age (Darst et al., 2019; Houtkooper et al., 2011; Johnson et al., 2018; Panyard et al., 2022; Puurunen et al., 2022; Sol et al., 2023; Wang et al., 2023). Because amino acids can be synthesized *de novo* or generated from proteolysis of either cellular or dietary protein, the physiological source of age-associated amino acids has not been identified, though several studies speculate that protein catabolism may vary by age (Lawton et al., 2008; Panyard et al., 2022). Measurement of ptmAAs on the other hand, which has only recently become more common in targeted metabolomic analysis, offers a clearer picture of the contributions from protein catabolism versus *de novo* synthesis (Darst et al., 2019; Mardinoglu et al., 2015). Here we found many ptmAAs associated with age, indicating that protein catabolism is an important aspect of age-related metabolome variation. Further support for this hypothesis was provided by the fact that the concentration of ptmAAs within the plasma of dogs was associated with BUN and creatinine, two biomarkers of amino acid catabolism **(Figure 3B**). Together, this evidence points to protein catabolism as a physiological nexus for age-associated variation in plasma metabolites among companion dogs.

We investigated several potential physiological influences on the abundance of ptmAAs in plasma, looking for those that might explain the age-association in particular. Diet can have large influences on the human serum metabolome (Bar et al., 2020). We found evidence that diet influences the abundance of two ptmAAs—S-methylcysteine and N-Ac-aspartate. The abundance of N-Ac-aspartate, which were age-associated, was lower in dogs whose primary diet was raw commercial food. Effects of pet-food processing on nutritional content is well studied and includes limiting the availability of amino acids from cooked food (Oba et al. 2022). S-methylcysteine was not age-associated and can form on internal cysteine residues, in contrast to the N-Ac-AAs, which are only formed on the N-terminus of protein. N-Ac-aspartate was the only N-Ac-AA negatively associated with age. However, among dogs whose primary diet type was dry kibble, N-Ac-aspartate was not age-associated. Therefore, it may be that the N-Ac-AAs, which were generally higher in older dogs, reflect a common physiological cause, somewhat distinct from that which gives rise to S-methylcysteine, and that the negative association of N-Ac-aspartate with age was simply explained by diet.

For there to be more ptmAAs in the plasma of old vs. young dogs, the rate of their addition to the plasma must be greater than their rate of removal or metabolism. Studies of digestive enzymes and digestibility in young and old dogs of several breeds, either in a >100-dog cohort, or in a controlled setting, indicate that older dogs more readily digest protein (Buddington et al., 2003; Weber et al., 2003). Thus, the elevated plasma ptmAAs we found in older dogs may be due in part to increased generation from dietary protein, all else being equal. In addition, we found that the lower rate of removal by glomerular filtration in the kidney in older dogs could explain their elevated ptmAAs. Glomerular filtration rate (GFR) is typically estimated as a function of the inverse of serum creatinine. While we do not estimate the GFR in this study, we found that approximately 50% of the effect of age on those ptmAAs that were higher in older dogs can be accounted for by variation in creatinine among the dogs (**Figure 4A**). Creatinine positively covaries with the abundance of ptmAAs generally (**Figure 3B**), suggesting that high creatinine, an indication of low GFR, leads to higher ptmAAs. This result is consistent with the association between GFR and N-Ac-ornithine in human blood <u>(Suhre et al. 2011)</u>. Therefore, the elevated ptmAAs in the plasma of older dogs appears to be due to lower rates of removal by glomerular filtration.

Another indicator of kidney function, uSG (McGlynn et al., 2023), associates with the ptmAA hydroxyproline, the only age-associated ptmAA that was not mediated by serum creatinine. That uSG itself does not associate with creatinine suggests that the reduced hydroxyproline, and the increased N-Ac-AAs, in older dogs may have independent physiological explanations. Urine specific gravity is a direct measure of the amount of solute removed to the urine, and is considered a measure of efficiency in urine concentration. Hydroxyproline can be metabolized into toxic byproducts, and so we would assume that the removal of hydroxyproline to the urine would be at its most efficient at high uSG (Belostotsky & Frishberg, 2022). The positive association between uSG and plasma hydroxyproline is therefore paradoxical. However, it is possible that uSG rises and falls directly or otherwise in response to hydroxyproline. Additionally, while maximal uSG is an indicator of renal function, uSG also varies in response to water consumption throughout the day and can also be affected by endocrine disease. Thus at any given point in time it may not be possible to determine whether or not a dog’s uSG represents its maximal renal concentrating ability and this may confound the ability to associate uSG with metabolite concentrations

Hydroxyproline is a major constituent of collagen and is reasoned to indicate tissue degradation, including muscle wasting, liver injury and fibrosis, or may reflect the level of dietary animal protein (Keiser et al., 1963). We failed to found an association between hydroxyproline and the primary diet component. An alternative explanation is that its age-related decline somehow reflects sarcopenia in older dogs. Sarcopenia, the degradation of endogenous muscle tissue with age, is a hallmark of aging in animals (Attaix et al., 2005; Fielding et al., 2011; Saini et al., 2009). Several studies have attempted to characterize serum or plasma metabolites associated with sarcopenia in humans. However, a consensus on plasma biomarkers of sarcopenia has not been reached, including an inconsistent association between plasma hydroxyproline and age-related muscle loss (Kameda et al., 2020, 2021; Pan et al., 2021). Here plasma hydroxyproline levels decline in older dogs, which is inconsistent with elevated levels of muscle and liver cell degradation as dogs age.

## Conclusion

The DAP is designed, in part, to develop a companion dog model of aging, which could provide major insights into healthy aging in one of the most variable species of mammal in terms of longevity, behavior, morphology and pathophysiology (Creevy et al., 2022). Conceptually, the metabolome provides us with a mechanistic bridge in the genotype-phenotype map (Harrison et al., 2020). As such, the metabolome can indicate the ways in which genotypic variation leads to the variation in longevity and healthspan that is exemplified by companion dogs. We use the age-associated plasma metabolome as a window into the physiological processes that vary with age in dogs and found that protein catabolism might provide insight into aging. The results presented here come with the important *caveat* that they represent patterns in a cross-sectional cohort. As the Dog Aging Project progresses, it will be important to examine longitudinal patterns, asking how metabolites change with age within individual animals as they age. In fact, given the demographic bias that may exist among the baseline Precision Cohort, as with cross-sectional studies of humans, we may be observing what healthy aging looks like among the older subjects in this study. Having identified age-associated change in about 39% of the aqueous metabolome, we have considerable leverage to detect environmental and genetic factors that influence the pace of aging in the longitudinal phases of the DAP, and to identify physiological processes that may respond to aging interventions and their effect on longevity and geriatric health.

## Materials and Methods

The Dog Aging Project (DAP) is a long-term longitudinal study of companion dogs in the United States. The project is designed to identify the genetic and environmental factors that influence age-related morbidity and mortality, and the mechanisms by which they do so. Dogs in DAP were recruited with the goal of retaining them for their lifetime. For the Precision Cohort dogs, beginning with the first year of enrollment and at one-year intervals thereafter, owners bring their dog to their primary care veterinary clinic for biospecimen collection. During this visit, clinical data were recorded, including the dog’s age, weight, sex and sterilization status, as well as the duration of fasting prior to blood collection (Prescott et al., *in prep*). Additionally, a veterinarian or veterinary technician collects hair, urine, a fecal sample, and blood samples, the last of which were separated into whole blood, plasma, serum, and peripheral blood mononuclear cells (Prescott et al., *in prep*). One aliquot of plasma was used for targeted aqueous LC-MS metabolomics.

### Blood Sampling, Plasma Extraction and Metabolite Extraction

For full details on the design and execution of dog owner contact and sampling in the Precision Cohort, see Prescott et al. (*in prep*). Briefly, blood samples in EDTA tubes, from either one draw of 20mL for dogs > 8kg, or two draws of 12mL each, six weeks apart, for dogs ≤8kg, were shipped to the Texas Veterinary Medical Diagnostic Laboratory. Along with sample appearance and other qualitative checks, the travel time and arrival temperature were recorded. For the samples measured here, the median travel time was 26.3h (range from 14.2 to 168.7h). The median arrival temperature was 18.8°C (ranged from 1.8 to 28.7°C). Plasma was extracted and transferred to 250μL aliquots in cryovials at the DAP Central Lab at Texas A&M University and stored frozen at −80°C until shipment to the University of Washington (UW). At UW, plasma samples were checked for hemolysis based on the Center for Disease Control and Prevention Hemolysis Reference Palette (CDCHRP, **Figure S4**). Metabolite extraction was performed at the UW Northwest Metabolomics Research Center (NW-MRC) in batches of up to 40 samples using a cold-methanol extraction protocol (Prescott et al., *in prep*) and stored at −80°C.

**Figure S4.**
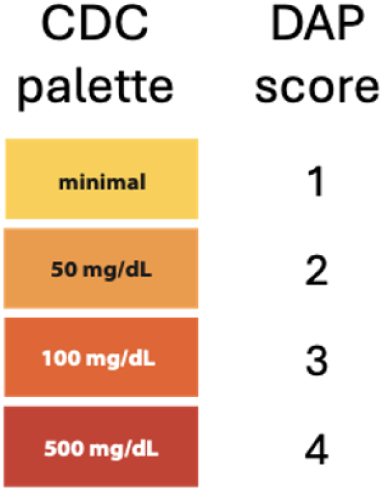
Hemolysis Reference Palette. The colorimetric reference palette used by the Dog Aging Project (DAP) to assess plasma sample hemolysis. Plasma samples in polypropylene microfuge tubes were compared to the Center for Disease Control and Prevention (CDC) palette and given a score from 1 to 4. Samples with a score of 4 were not analyzed.

Prior to LC-MS, samples were reconstituted in 500 µL HILIC solvent containing ^13^C_2_-tyrosine, ^13^C_3_-lactate and 10mM ammonium acetate in 5% methanol and 0.3% acetic acid. To track secular trends in LC-MS detection that occur during the run, multiple replicates of two different control samples were included in each LC-MS experiment. These include a reference dog plasma sample from the Cornell Veterinary Biobank, referred to as QC(S), and an NW-MRC human plasma control sample. Each of these was run first, then again, interspersed across the experiment, between every ten DAP samples, and at the end of the run. LC-MS peaks were integrated to give metabolite count data, which were sent from the NW-MRC back to the Promislow Lab. Raw LC peak and MS spectral data are stored on servers at the NW-MRC.

The chromatography separations were performed on a duplex-LC system composed of two Shimadzu UPLC pumps (Shimadzu Corp., Columbia, MD), Agilent 1290 temperature-controlled column compartment (Agilent Technologies, Santa Clara, CA) and CTC Analytics PAL HTC-xt temperature-controlled auto-sampler (LEAP Technologies, Morrisville, NC). The LC modules were controlled by Analyst 1.7.2 software (AB Sciex, Toronto, ON, Canada). Each sample was injected twice, 10 µL for analysis using negative (NEG) ionization mode and 5 µL for analysis using positive (POS) ionization mode. Both chromatographic separations were performed in HILIC mode on two XBridge BEH Amide columns (150 x 2.1mm, 2.5µm particle size, Waters Corporation, Milford, MA, Part No. 186009930) connected in parallel. While one column was performing the separation, the other column was reconditioned in preparation for the next injection. The flow rate was 0.300 mL/min, auto-sampler temperature was kept at 4°C, the column compartment was set at 45°C, and total separation time for both ionization modes was 18 min (total analysis time per sample was 36 min). The mobile phase in POS mode was composed of Solvents A (10mM ammonium acetate in 95% H_2_O/3% acetonitrile/2% methanol + 0.2% acetic acid) and B (10mM ammonium acetate in 93% acetonitrile/5% H_2_O/2% methanol + 0.2% acetic acid). The gradient conditions for POS mode separation are shown in **Table 1**.

**Table 1:** LC Gradient Conditions for POS Mode Chromatography.

| Time Segment, min. | Solvent A, % | Solvent B, % |
| --- | --- | --- |
| 0 - 3 | 5 | 95 |
| 3 - 8 | from 5 to 50 | from 95 to 50 |
| 8 - 12 | 50 | 50 |
| 12 - 13 | from 50 to 5 | from 50 to 95 |
| 13 - 18 | 5 | 95 |

**Table 2:** LC Gradient Conditions for NEG Mode Chromatography.

| Time Segment, min. | Solvent A, % | Solvent B, % |
| --- | --- | --- |
| 0 – 1.5 | 5 | 95 |
| 1.5 - 6 | from 5 to 30 | from 95 to 70 |
| 6 - 10 | 30 | 70 |
| 10 - 12 | from 30 to 55 | from 70 to 45 |
| 12 - 14 | 55 | 45 |
| 14 - 15 | from 55 to 5 | from 45 to 95 |
| 15-18 | 5 | 95 |

### Data Transformation and Technical Covariates

The LC-MS data were composed of peak intensity values for 361 metabolites among a total of 1346 samples. This includes 920 baseline Precision Cohort samples, with the remaining samples belonging to other DAP cohorts. These data were collected over the course of five LC-MS runs. In each run, between 73 and 600 samples were run in the same order in which they were processed during metabolite extraction. This design maximizes the statistical power to detect and remove batch and LC-MS run-order effects. We normalized and pre-processed all LC-MS data together. We removed 224 metabolites across the study due to missingness in >10% of all samples. The level of hemolysis was associated with the abundance of some metabolites, and 72 samples were removed due to hemolysis exceeding 500 mg/dL (CDCHRP score=4, **Figure S6**). The remaining data were log_e_-transformed, and mean-centered by sample to account for sample-to-sample variation in metabolite abundance.

Both metabolite extraction (batch) and LC-MS runs generate secular variation in metabolite data. Such variation could be due to undefined chemical reactivity, sample matrix effects, ion suppression (interference), etc., and thus differs by metabolite. In these cases, the LC-MS peak corresponding to an affected metabolite drifts with the order in which a sample was processed within a batch. There is also the potential for peak area to vary with the order of samples within an LC-MS run. To correct both for main effects of batch, and for run-order effects, we take the residuals (*e*) of the regression model:

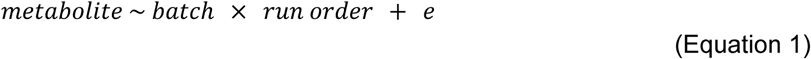

This results in peak areas that are both normalized, and that correct for effect of run order. To correct for variation in dynamic range among metabolites within experiments, we scaled each metabolite to unit variance by batch. After these procedures, there were an average of 6 missing values per metabolite, with at most 125 missing (9.8%). These missing values were then imputed by 10-nearest neighbor mean-imputation. Effects of the remaining technical covariates, travel time, hemolysis and arrival temperature on metabolite abundance were corrected for by linear regression. The data processing and normalization resulted in 137 normalized metabolites measured in 865 dogs in the baseline Precision Cohort.

### Breed Ancestry and Genetic Relatedness

Each dog of the Precision Cohort has had low-pass whole genome sequencing (Sexton et al., *in prep* 2024). Briefly, reads were aligned to the CanFam3.1 reference genome assembly (NCBI accession GCF_000000145.2) and imputation using a panel of reference haplotypes including >34M SNPs and >11M indels shared by 109 modern dog breeds, 3 village dog populations and North American and European wolf populations (Sexton et al., *in prep* 2024). SNPs with a minor allele frequency greater than 1% and genotype call rates greater than 95% were retained. Here we used the genetic data in two ways—first, to determine breed by genetic ancestry and evaluate the effect of such breed information on metabolome profiles, and second, to control for relatedness among all dogs regardless of breed. For breed ancestry, we estimated the proportion of genetic ancestry in each dog genome using publicly available genotype data from 109 modern breeds, village dog populations from three regions, and two wolf populations. Ancestry proportions for each dog were then estimated using ADMIXTURE on genotype data from 115,427 biallelic SNPs. Where applicable here, purebred was defined as any dog with ≥85% ancestry assigned to a single breed (Morrill et al., 2022). To estimate relatedness, the variance-standardized GRM was calculated with autosomal markers in PLINK2 using the default settings in the make-rel function (Chang et al., 2015).

### Variable Selection and Normalization

Of the data collected on Precision Cohort dogs, age and weight were square root transformed, and sex (male and female) and sterilization status (intact or sterile) were coded as factors (i.e., 0 and 1). The duration of fasting prior to blood collection was rounded to the nearest hour. Based on the American Animal Hospital Association’s Canine Life Stage Guidelines, Precision Cohort dogs were classified into one of five age categories: puppy (<1 year), adolescent (1-3 years), young adult (3-7 years), mature adult (7-11 years), and senior (11+ years) (Prescott et al *in prep* 2024).

Complete Blood Count and blood chemistry (CBC/Chem) data were acquired at the Texas Veterinary Medical Diagnostic Laboratory, from samples in the DAP biospecimen kits. CBC measures were taken from blood samples in EDTA tubes on an Advia 120 Hematology System (Siemens Medical Solutions, Malvern, PA) from 784 of the Precision Cohort dogs that had metabolome profiles. Parallel blood chemistry profiles were performed with serum extracted from an accompanying additive-free tube, and run on a DxC700AU Chemistry Analyzer (Beckman Coulter, Brea, CA). Together the raw data consist of 45 CBC and 42 chemistry measures. The 36 dogs without CBC/Chem data were removed from further analysis. Of the 45 CBC traits, we removed five invariant measures, 36 that had >10 missing values, and any that were relative measures when an absolute measure was available. Among the remaining 39 variables, ten of the numerical CBC variables were non-normal, including counts of band cells, neutrophils, lymphocytes, monocytes, eosinophils, basophils and reticulocytes, red cell distribution width, mean platelet volume, and plateletcrit (Shapiro-Wilk statistic <0.96), and were log_e_-transfomed. There was an average of one missing entry among each of the variables at this point (maximum missiness was five). All missing data were imputed by 10-nearest neighbor mean imputation, which gave 38 complete and normalized CBC/Chem variables in 784 dogs (**Table S2**). The effect of blood sample travel time and arrival temperature on each CBC trait were removed by linear regression. The adjusted 17 CBC variables were used as covariates in mixed models.

Urinalysis was performed with approximately 3 mL of urine on 738 of these 784 dogs. The uSG was calculated by refractometer; chemical analysis was performed with Multistix 10 SG Urine Test Strips (Siemens Medical Solutions, Malvern, PA); and microscopy was performed manually (Prescott et al., in prep). After removing two invariant urinalysis variables, there were two numeric variables: uSG and pH, and 12 categorical variables. Eight of the categorical variables: protein, white blood cells, red blood cells, squamous cells, urothelial cells, bilirubin, fat and blood, were ordered semi-quantitatively. For example, urothelial and other cell counts were coded ‘None Observed’ < ‘Rare’ < ‘0-3’ < ‘3-6’ < ‘6-10’ < ‘10-20’ < ‘20-40’ < ‘Too Numerous to Count’. The remaining categorical variables, including urine color, transparency, crystals and casts, were not evaluated. This gave 10 clinical urinalysis variables (**Table S2**). As covariates, the ordered categories were converted to integers and, along with the continuous numeric variables, were mean-centered and scaled to unit variance prior to model fitting with each as a fixed effect, using the mixed model described below (Equation 2).

**Table S2.**
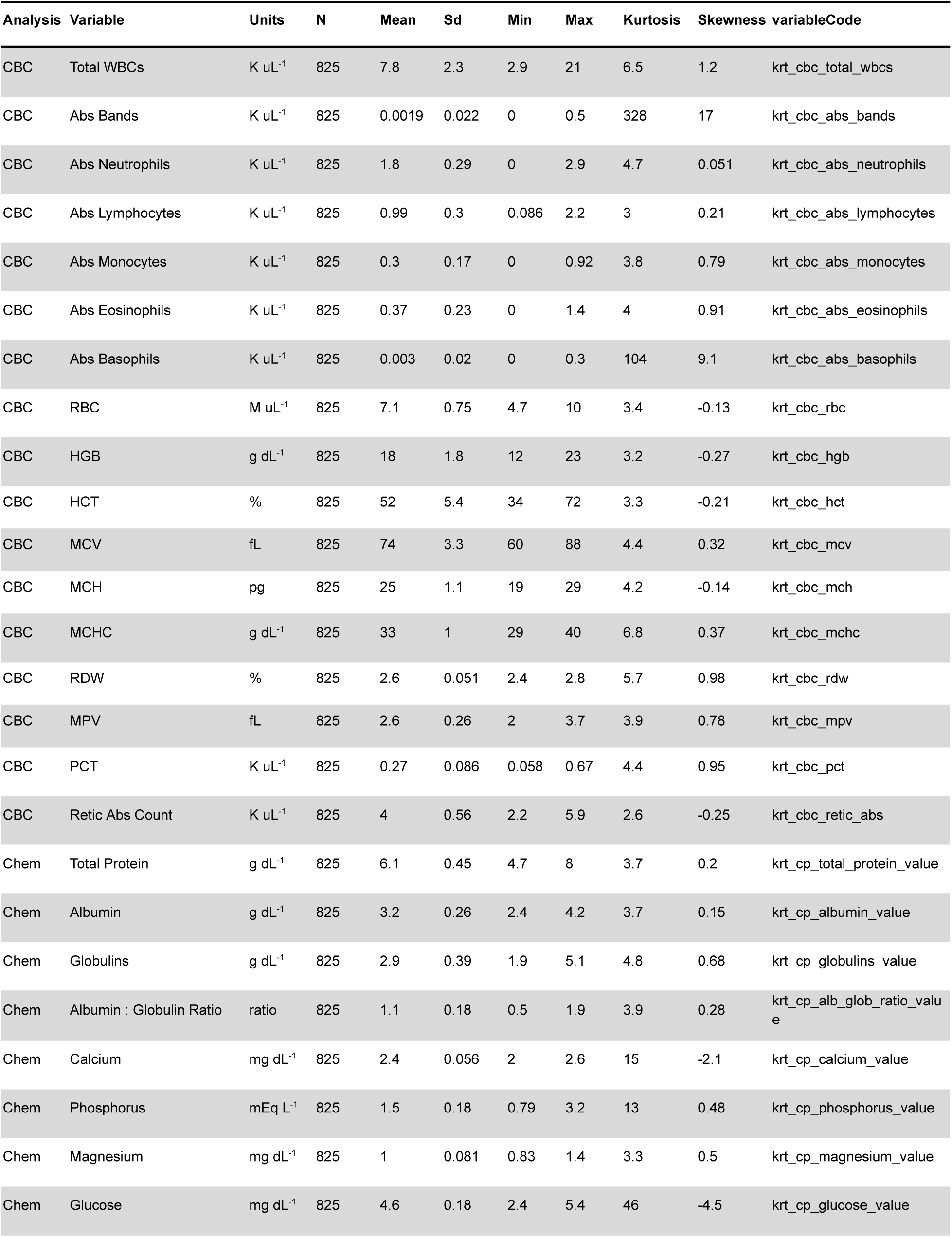
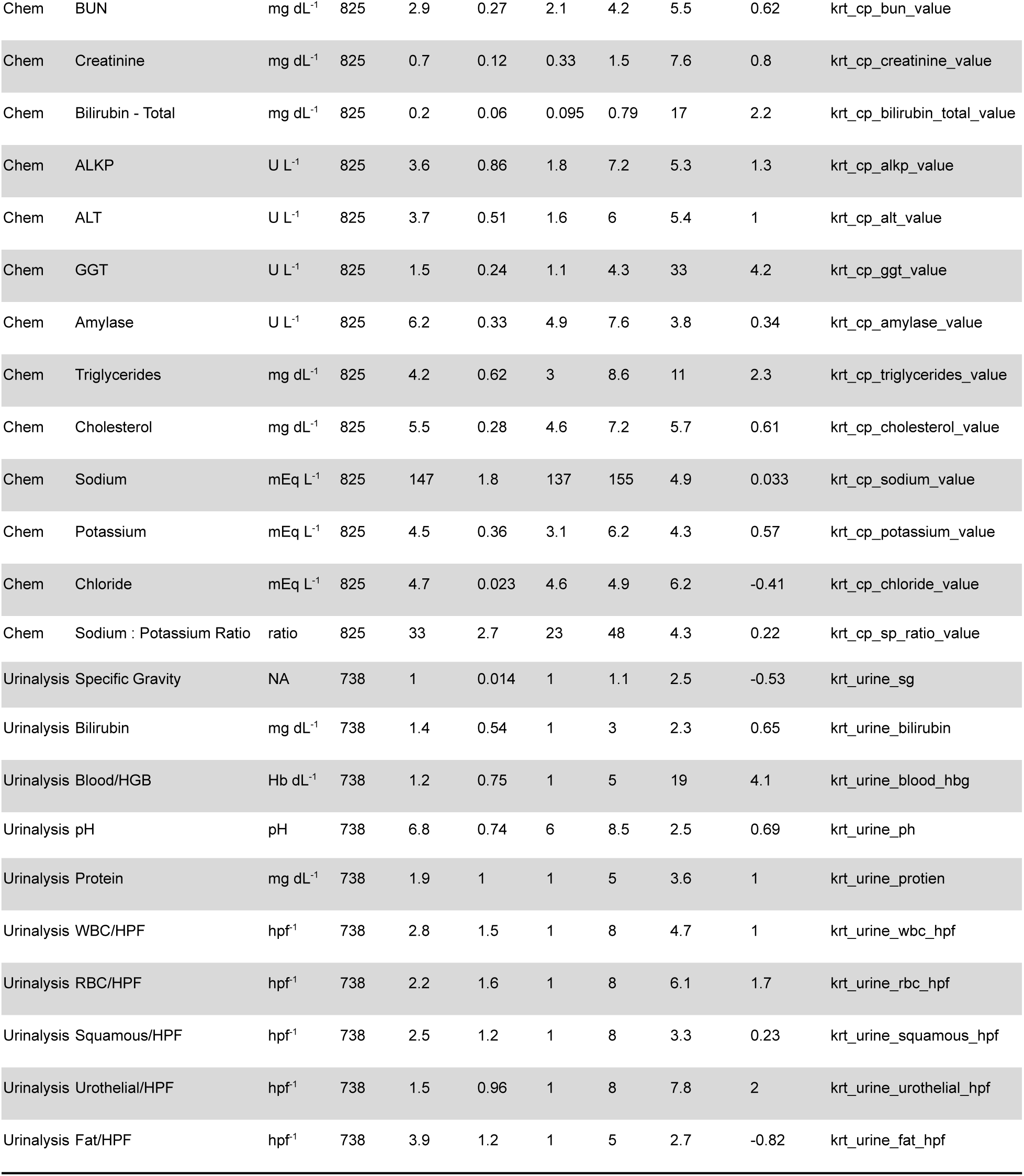
Summary of Complete Blood Count, Serum Chemistry and Urinalysis Variables. The names, units and summary statistics: N=number of dogs with available data, SD=standard deviation, and measures of skewness and kurtosis, for variables selected for analysis based on completeness and other factors (Methods). The variable names in the meta data are given (variableCodes). There were 17 variables selected from the complete blood count (CBC) analysis, (variableCodes contain ‘krt_cbc’), 21 serum chemistry variables (Chem, variableCodes contain ‘krt_cp’), and 10 variables from the Urinalysis (variableCodes contain ‘krt_urine’).

### Principal components analysis

Principal component analysis was performed on normalized metabolome data from 784 dogs for which we also had CBC data, with additional scaling by metabolite. We used the Tracy-Widom test in the *AssocTest* package to identify the first 23 PCs that describe significantly non-random variation (α = 0.05). Type III ANCOVA was used to estimate the variance of each of the first 23 PCs that could be explained by effects of each covariate.

### Linear Mixed Model

The fixed effects (β) of *X*, the design matrix of the covariates: age, weight, sex, sterilization status, the duration of fasting prior to blood collection, the 17 CBC traits, and the interactions between age and weight, and sex and sterilization status, were fit simultaneously on each metabolite (*y*), along with the random effects 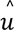 (best linear unbiased predictions, *BLUPs*) of the covariance in the GRM.

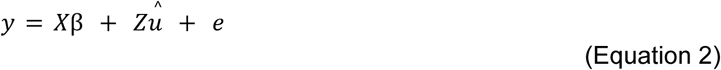

The mixed model was fit by maximum likelihood in the *EMMREML* package (Akdemir & Okeke, 2015). We tested for significance (β>0) of fixed effects within the emmreml function, and the P values were corrected for multiple comparisons by the false discovery rate method (FDR, (Benjamini & Hochberg, 1995)). When comparing models with and without the GRM, we fit an identity matrix (diagonals =1, off-diagonals =0) in place of the GRM.

### Adjustment for Fixed and Random Effects

When assessing the correlation among the ptmAAs, creatinine, BUN, and the unmodified amino acids, we removed the effects of the fixed covariates from the metabolite values (*y*) by subtracting both fixed effects from Equation 2 (the best linear unbiased estimators, *BLUEs*), and random effects (*BLUPs*) from Equation 2 using equation 3:

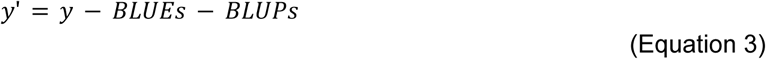

where BLUEs were derived by multiplying the design matrix (*X*) by the matrix of fixed effects (β), which, when subtracted from *y*, give the fully-adjusted values (*y*’).

For metabolite covariation analysis, Pearson r was calculated on pairs of fully-adjusted metabolite values (*y*’) and the measures of similarity among the resulting r among all pairs were clustered by the unweighted pair group method with arithmetic mean UPGMA in R.

### Mediation analysis

We performed mediation analysis with the *mediation* R package (Tingley et al., 2014), testing for the causal mediation effect (γ) of a mediator (*M*) on the effect of age (β) on a metabolite. Mediation was estimated in linear models with the same fixed-effects covariates (*X*) used in the mixed model (Equation 2), without the random effect of the GRM (Equation 4):

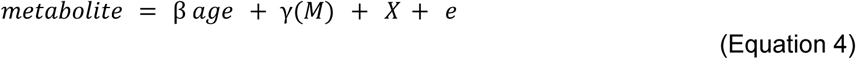

The causal mediation effect (**Figure S5)** was estimated and tested for γ>0 using up to 10^6^ bootstrap randomizations of age. The proportion mediated is given by dividing γ by the effect of age on a metabolite without the mediator (**Figure S5**). We tested the sensitivity of γ to unmeasured confounding among the predictors by sensitivity analysis, where correlation (ρ) between the residual effects of the mediator and outcome variables was artificially introduced to estimate the ρ at which γ=0 (Imai et al., 2010). None of the mediation models with γ>0 at FDR≤5% were sensitive.

**Figure S5.**
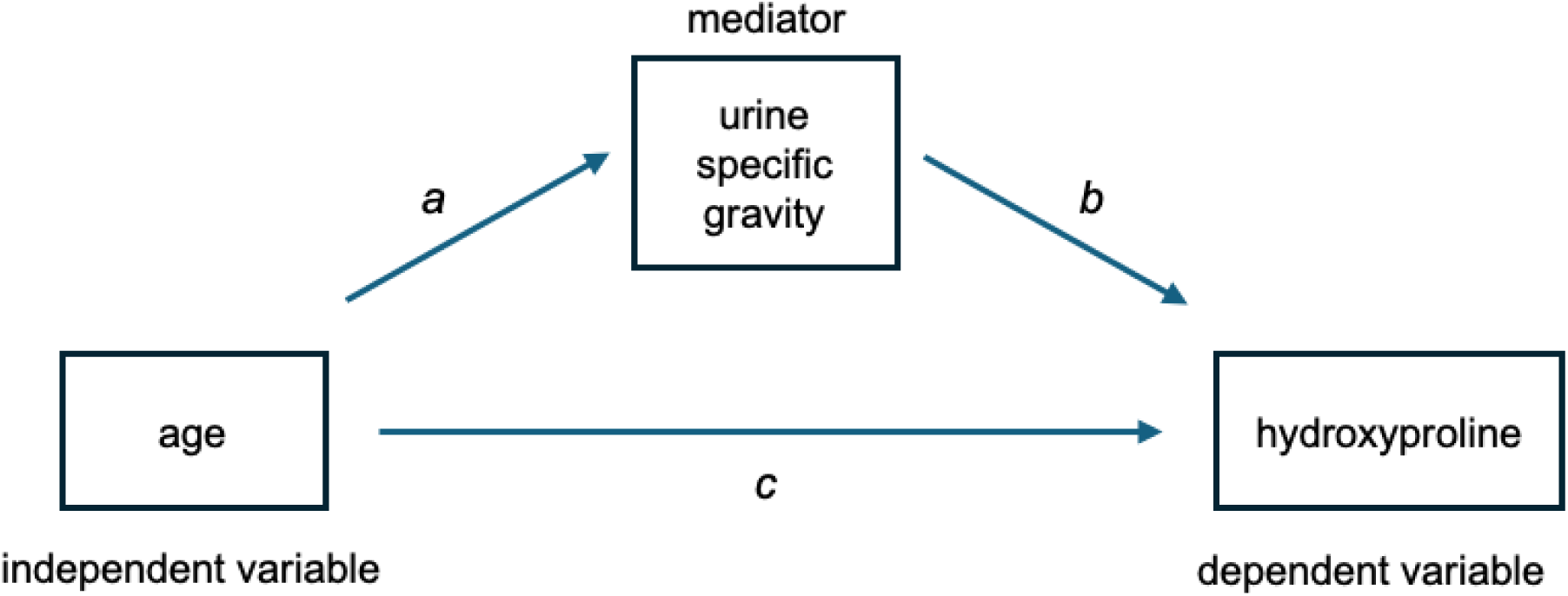
Path representation of causal mediation models. A path representation of a causal mediation model using, as an example, the hypothetical effect of age on hydroxyproline, with urine specific gravity as a potential mediator. The mediation model (path a-b), was compared to a model without urine specific gravity (path c). The total effect was path c, without considering path a-b. To detect causal mediation, after satisfying the assumption that path a was significant, the direct effect (c|b) of urine specific gravity was the effect of path c given path b. The mediation effect was the total effect minus the direct effect, and its significance (γ>0) was tested by bootstrap resampling age in the model (Methods). The proportion mediated was γ divided by the total effect.

## Acknowledgements

The authors thank Dog Aging Project participants, their dogs, and community veterinarians for their important contributions.

## Data and Code Availability

Dog Aging Project data are available on the TERRA platform at the Broad Institute of MIT and Harvard (https://app.terra.bio/). Code for this study will be made available on GitHub: https://github.com/ben6uw/DAPmetabolome upon publication.

